# Modifying the aggregation state may improve the activity of Ozempic

**DOI:** 10.1101/2024.11.19.624403

**Authors:** Alain Bolaño Alvarez, Kristian B. Arvesen, Kasper F. Hjuler, Peter Bjerring, Steffen B. Petersen

**Affiliations:** Department of Dermatology and Venereology, Aalborg University Hospital, Hobrovej 18-20, Aalborg, Denmark

## Abstract

We here report a study of aggregation of Semaglutide at different temperatures, using resonance light scattering (RLS), fluorescence polarization and back-scattering techniques. Fluorescence emission spectra were obtained by exciting the samples at 275 nm and 295 nm, revealing a peak emission at 600 nm associated with the aggregation process. The size of the aggregates is around 100 nm according to back-scattering measurements. Two distinct thermal transitions were observed by RLS: the first melting point (Tm1) at 30°C and the second (Tm_2_) at 91°C, indicating changes in aggregation state. The fluorescence polarization revealed a fast rotational dynamics of the aggregates at Tm_2_, leading to greater depolarization of the emitted light. The structural organization of the Ozempic aggregates was studied using two dyes, Laurdan for lipid components and 1,8-ANS for protein component (GLP1). Thus, revealing a stable PEG-lipid core which hold the GLP1, increasing their exposition to the solvent. An enhanced FRET event inside the aggregates in presence of Fe^2+^ and Fe^3+^ was observed. We conclude that the PEG-lipid core plays a significant role in the aggregates structure stability, being a key to improve the biological activity of Ozempic. This methodology can be used to study similar aggregation constructs in the pharmaceutical industry.

## Introduction

The aggregation behavior of protein-based biopharmaceutical compounds, particularly under thermal stresses, is crucial for understanding and optimizing their stability, efficacy, therapeutic performance, storage, and delivery [1, 2]. Thus, in modern pharmaceutical formulation the aggregation can provide benefits to improve aspect such as the therapeutic performance and stability. It is the case of Semaglutide a product from Novo Nordic, Denmark [3], based on semaglutide peptide as active component, which has GLP-1 receptor agonist activity, this medication is widely prescribed for managing type 2 diabetes [4]. The chemical structure of active principle in Ozempic comprising of two poly(ethylene) glycol (PEG) units incorporated as Fmoc-Lys(Dde)-OH to the side chain of Lys^26^ in the peptide sequence, a γ-glutamic acid to bind the PEG units to a saturated fatty di-acid C_18_-OH [5]. In this pharmaceutical construct the incorporation of both PEG and fatty di-acid C_18_-OH in semaglutide peptide, plays a significant role in enhancing its therapeutic efficacy and pharmacokinetic profile of Ozempic [6, 7].

This study demonstrates the presence of aggregates in Ozempic formulation using polarization fluorescence spectroscopy and resonance light scattering (RLS). Also, we described molecular changes within its PEG-lipid core and the protein component across a range of temperatures claiming its potential impacts on bioavailability and function.

By identifying two critical melting points, the study provides insight into the molecular organization and stability of Ozempic. Furthermore, fluorescence resonance energy transfer (FRET) experiments reveal structural modifications induced by Fe^2+^ and Fe^3+^ ions, suggesting its importance for optimizing Ozempic therapeutic efficacy simply by molecular reorganization. These findings offer a new framework for enhancing biopharmaceutical formulations by modulating thermal aggregation and structural accessibility in complex drug systems.

## Results and Discussion

### Resonance light scattering and fluorescence polarization to define the aggregation state in Ozempic

Fluorescence emission spectra were measured by exciting the samples at 275 nm and 295 nm. A maximum emission at 600 nm was observed. This peak in the emission spectrum is closely associated with the aggregation process (**Figure 1A, green arrow**), which was further investigated through the determination of the G-factor using fluorescence polarization (**Figure 1C**).

**Figure 1:**
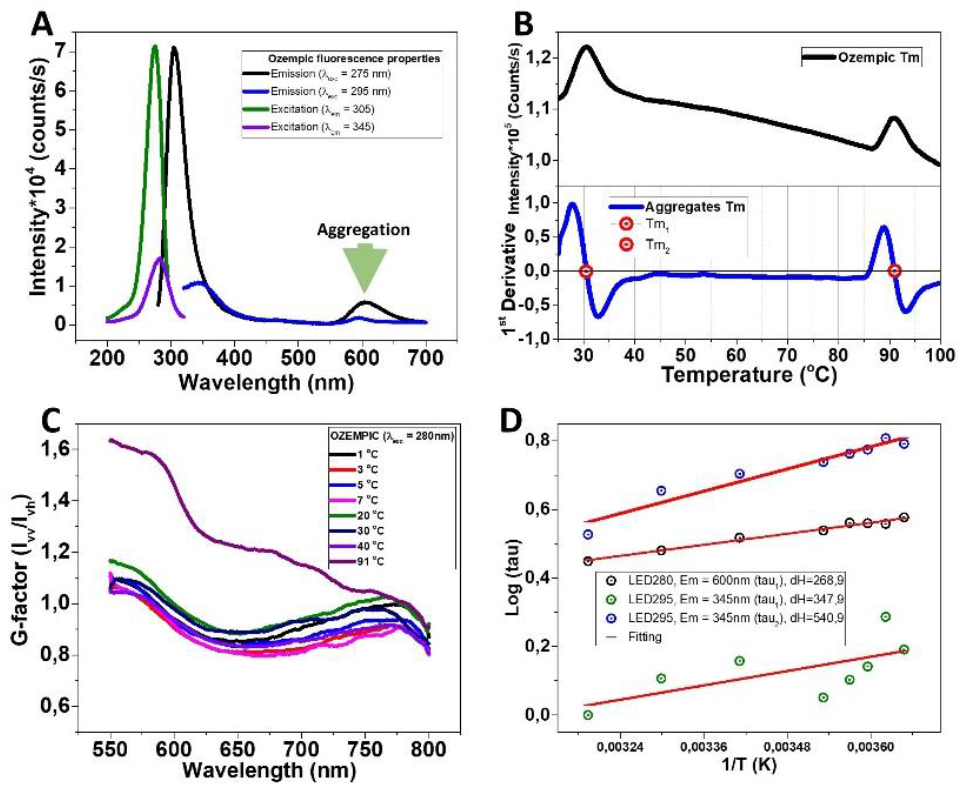
Melting point and fluorescence polarization to find the aggregation in Ozempic. **A:** Resonance light scattering to determine two melting points in Ozempic (Tm_1_ = 30°C and Tm_2_ = 91°C). **B-C:** G-factor determination to define the rotating states of the Ozempic aggregates under different temperatures values. **D:** Temperature influence in the folding event of the aggregates.

The melting points of the Ozempic aggregation were determined using resonance light scattering (RLS) [8]. Two distinct thermal transitions were observed: the first melting point (Tm1) occurred at 30°C, and the second melting point (Tm_2_) occurred at 91°C. These transitions correspond to changes in the aggregation state of the Ozempic molecules in solution (**Figure 1B**).

To further probe the relationship between the aggregation state and thermal transitions, the G-factor was determined through fluorescence polarization experiments. These measurements were performed at different temperature values, specifically focusing on the wavelength range of 550 nm to 800 nm, where an emission peak was detected at 600 nm, which we relate with the aggregation emission. These measurements were achieved by placing polarizers in both the excitation and emission light paths of the fluorometer, allowing precise control over the polarization of the light. This depolarization effect can be explained as follows: when a fluorophore population is excited with vertically polarized light, the emitted light retains some degree of that polarization depending on the rotational speed of the molecules in solution. Faster rotational motion leads to greater depolarization emission, while slower motion retains more of the polarization emission [9].

The G-factor showed a marked increase at Tm_2_ (91°C), indicating a significant change in the rotational dynamics of the aggregates. At this temperature, when the aggregates were excited with vertically polarized light, the emitted light was notably more depolarized. This suggests that the aggregates were rotating more rapidly at Tm_2_ compared to Tm_1_ (30°C), where the polarization retention was higher, indicating slower rotational motion (**Figure 1C**).

The results at Tm_2_ demonstrate that the aggregates undergo faster rotational motion at higher temperatures, causing the emitted light to become more depolarized.

To further explore the relationship between PEG-Lipid component aggregation, protein component structural change, and temperature, we conducted lifetime (τ) measurements. Emission at 345 nm (excitation at 295 nm) provided almost two distinct lifetimes, τ_1_ and τ_2_, which correspond to the tryptophan structural transition in the excited state [10] of Ozempic protein component. Additionally, emission at 600 nm (excitation at 280 nm) revealed a single lifetime, τ_1_, which is attributed to the Ozempic aggregates. Notably, the lifetime associated with the aggregates displayed longer τ values, a characteristic that is typically linked to the aggregation process.

We plotted the logarithm of the lifetime (log(τ) versus the inverse of temperature (1/T, in Kelvin) to examine the enthalpy changes. The linear behavior of these plots indicates a thermally activated process, where the slopes of the fitted lines correspond to changes in enthalpy (**Figure 1D**). For emission at 600 nm (τ_1_, corresponding to aggregates), the enthalpy change was determined to be 268.9 kJ/mol. For emission at 345 nm (τ_1_, corresponding to S_0 →_1L_b_ tryptophan transition), the enthalpy change was 347.9 kJ/mol. For τ_2_ (S_0 →_ 1L_a_ tryptophan transition) at 345 nm, the enthalpy change was 540.9 kJ/mol.

Since all the enthalpy values are positive, this indicates that the PEG-Lipid component aggregation and protein component structural change events are endothermic processes. The enthalpy changes correspond to the energy required for the thermal structural change of the protein component and the rearrangement of the PEG-Lipid aggregates. These findings suggest that the aggregation of Ozempic is a thermally influenced process, consistent with the observed melting points and fluorescence polarization results (**Figure B and C**).

### Fluorescence spectroscopy analysis to deduce aggregation structural organization

To elucidate the structural organization of Ozempic, we employed fluorescence spectroscopy using two fluorescent dyes—Laurdan, which inserts into lipid components [11, 12], and 1,8-ANS, which interacts with protein components [13, 14]. Additionally, we assessed changes in the polyethylene glycol 200 (PEG200) component by measuring the fluorescence of Ozempic in the presence of Fe^2+^ ions (**Figure 2**). Laurdan, a lipid-sensitive dye to label fluid lipid phase [11], was used to investigate the organization of the lipid component of Ozempic aggregates. The excitation wavelength was set at 385 nm, and fluorescence emission was measured after incubation at different time points: 1 hour, 2 hours, and overnight at room temperature.

**Figure 2:**
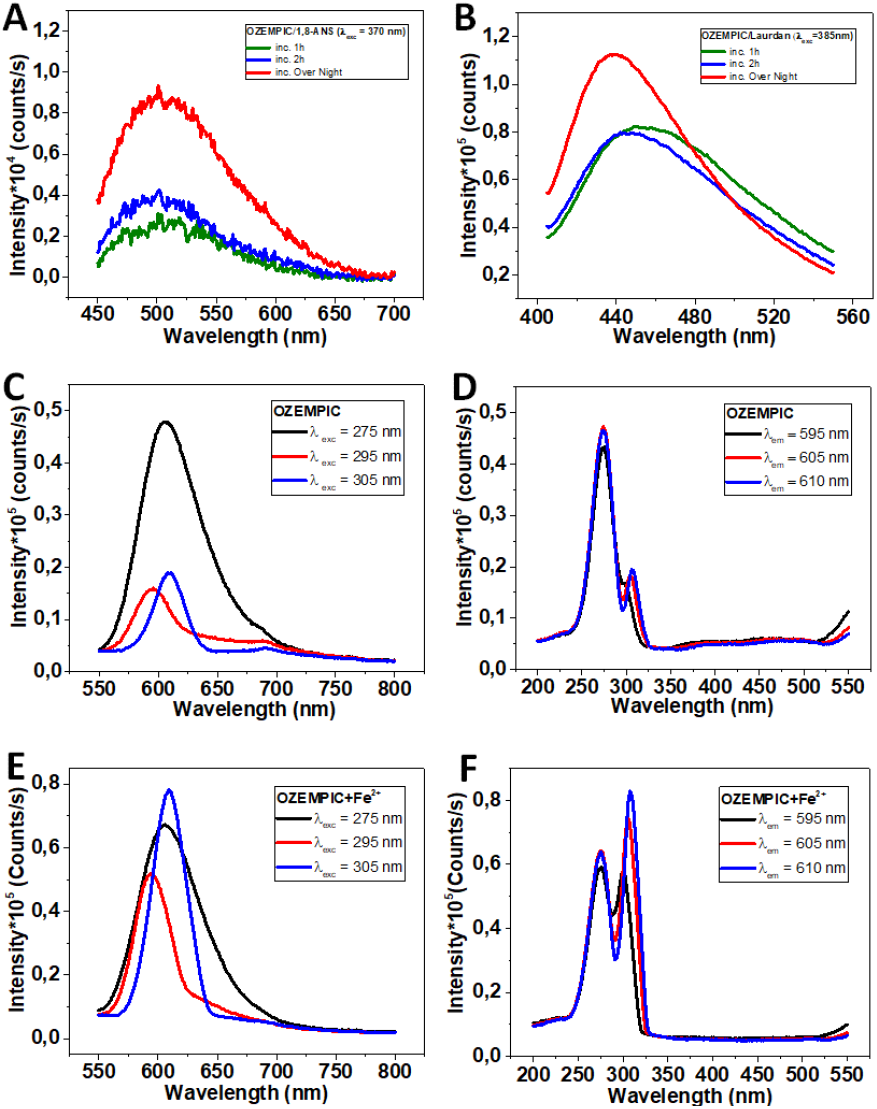
Molecular arrangements of the aggregates (Ozempic). **A:** Fluorescence emission of Ozempic in presence of 1,8-ANS. **B:** Fluorescence emission of Ozempic in presence of Laurdan. **C-D:** Fluorescence emission and excitation properties of Ozempic aggregation. **E-F:** Fluorescence emission and excitation properties of Ozempic aggregation in presence of Fe^2+^.

After 1 and 2 hours of incubation, no significant emission was observed, indicating minimal Laurdan insertion into the lipid component at these early time points. However, after overnight incubation, the emission at 440 nm increased, suggesting that the lipid tails had reorganized into a compact, ordered state within the aggregates. We suggest that this ordered phase arises with the incubation time and likely hinders Laurdan insertion during shorter incubation periods but allows for better insertion with longer incubation times. Indicating that the lipid component as part of the aggregates form a compact packed, which becomes more accessible to Laurdan with the incubation times.

We also used 1,8-ANS, a dye that specifically binds to protein regions, this fluorescence dye allows to study the interaction with the protein component of Ozempic. The excitation wavelength was set to 370 nm, and the emission was measured at 1 hour, 2 hours, and overnight.

Emission at 500 nm was detected after 1 hour, 2 hours, and overnight incubation, with gradual improvement in fluorescence intensity over time (**Figure 2B**).

In both cases, the lipid component and the protein portion also becomes progressively more accessible over incubation time. This supports the hypothesis that the protein component is part of the core structure within the aggregates, rendering it less accessible to 1,8-ANS in the early stages of incubation but allowing more interaction as the aggregates become more stable and ordered at the stored temperature.

It is well known that PEG has many advantages in pharmaceutical applications due its ability to form complex metal cations such as Fe^2+^ by interacting in the with the oxygen and hydroxyl group [15, 16]. Those PEG metal ions interactions allow determine whether the PEG200 is involved in the aggregation state, by studying the fluorescence properties of Ozempic in presence of Fe^2+^. We recorded both emission and excitation spectra and significant changes were observed immediately after the addition of Fe^2+^ ions, indicating a rapid interaction between Fe^2+^ and PEG200 (**Figure 2C-F**).

The immediate changes observed in both the excitation and emission spectra suggest that the PEG200 component of Ozempic is highly accessible to Fe^2+^ ions (**Figure 2E-F**). This rapid interaction highlights a key difference compared to the lipid and protein components, which require longer incubation times for detectable fluorescence changes with their respective dyes. Thus, we can conclude that the PEG component is located on the surface of the Ozempic aggregates, forming a kind of shield around the aggregates core, which consists of the lipid and protein components. At the same time, this shield increases the solubility of the aggregates and enhances the protein exposure on the aggregates surface to improve the Ozempic activity.

### FRET reveals a structural change in Ozempic influenced by Fe^3+^

In this study we have determined the overlapping integral in the region of fluorophore donor (Tyr) and acceptor (PEG-Lipid) involved in the energy transfer event (**Figure 3A**). This determination provides how the FRET efficiency changes indirectly in presence of Fe^3+^ and Fe^2+^ (**Figure S1**). The efficiency of this energy transfer depends on several factors, including the overlap of the emission of the donor spectrum and the absorption acceptor spectrum (overlapping integral, J(λ)), as well as a critical distance value between donor and acceptor in the range of 1-10 nm for effective FRET [17, 18].

**Figure 3:**
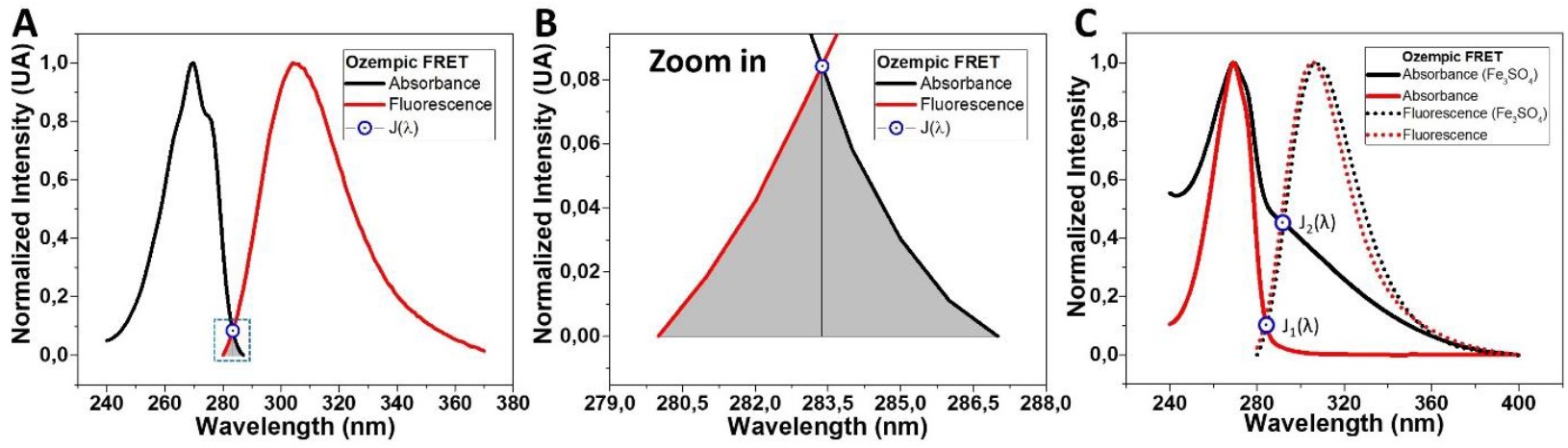
FRET in Ozempic molecular structure. **A:** Absorbance and emission spectra of Ozempic (λ_exc_ = 275 nm). **B:** Zoom in of the overlapping integral J(λ) in the region of donor (Tyr) emission and acceptor (PEG-Lipid aggregates) absorbance. **C:** Change in the energy transfer in presence of Fe^3+^ by influencing in donor-acceptor separation distance and their orientation.

In our study the low FRET efficiency was observed in a poor spectral overlap between the Tyrosine emission and PEG-Lipid absorbance, suggesting suboptimal distance and orientation for efficient FRET (**Figure 1A-B**). We have improved the FRET efficiency in presence of Fe^3+^ to modify the molecular structure of the aggregates, influencing both the distance and orientation of Tyrosine on the PEG-Lipid holder aggregates (**Figure 1C**). This change in J(λ) upon the introduction of Fe^3+^ indicates that Fe^3+^ interacts with the PEG-Lipid aggregates by coordination binding [19] in the Ozempic molecular assembly.

We suggest the presence of metal ions in the formulation might enhance the activity of Ozempic by changing the structure of the aggregates, possibly improving the arrangement of the protein component on the PEG-Lipid aggregates surface. Thus, this study provides a tool to study how optimizing the therapeutic function of Ozempic in biological contexts where FRET-related mechanisms play a role in similar molecular assembly.

## Conclusion

In this study, we characterized the aggregation and thermal stability of Ozempic using RLS and fluorescence polarization. Thus, two distinct melting points (Tm_1_ at 30°C and Tm_2_ at 91°C) were detected reflecting sensitivity to thermal transitions, which influence the rotational dynamics of the aggregation state in Ozempic active principal.

The aggregation structure where a central core based on PEG-lipid was deducted employing two fluorescent dyes, Laurdan and 1,8-ANS.

Additionally, the presence of Fe^2+^ and Fe^3+^ ions demonstrated FRET enhancement inducing a molecular rearrangements of the aggregates. It could be potentially considered in modern aggregation-based biopharmaceutical where a core PEG-lipid hold the active protein component as GLP1.

Our results highlight the importance of thermal modulation and controlled metal ion interaction to optimize drug delivery, solubility and stability in complex pharmaceutical construct.

